# Theta phase coordinated memory reactivation reoccurs in a slow-oscillatory rhythm during NREM sleep

**DOI:** 10.1101/202143

**Authors:** Thomas Schreiner, Christian F. Doeller, Ole Jensen, Björn Rasch, Tobias Staudigl

## Abstract

It has been proposed that sleep’s contribution to memory consolidation is to reactivate prior encoded information. To elucidate the neural mechanisms carrying reactivation-related mnemonic information, we investigated whether content-specific memory signatures associated with memory reactivation during wakefulness reoccur during subsequent sleep. We show that theta oscillations orchestrate the reactivation of memories, irrespective of the physiological state. Reactivation patterns during sleep autonomously re-emerged at a rate of 1 Hz, indicating a coordination by slow oscillatory activity.

---

The memory function of sleep relies on the reactivation of newly acquired information during non rapid-eye movement (NREM) sleep^1^. Rodent studies have consistently shown hippocampal reactivation of previous learning experiences during sleep^2^, while studies in humans have provided first hints indicating similar processes^3,4^. Furthermore, triggering reactivation processes during sleep by re-exposure to associated memory cues (targeted memory reactivation, ‘TMR’) has been shown to improve memory consolidation^5^.

However, the neural mechanisms coordinating reactivation-related mnemonic information are essentially unknown. Here, we investigated whether memory reactivation during wakefulness and sleep share oscillatory patterns that carry memory-representation specific information, using electroencephalography (EEG) and multivariate analysis methods.

Building on previous findings^6^, we hypothesized that low-frequency oscillatory phase conveys a representation (i.e. content)-specific temporal code. Using representational similarity analysis (RSA) to reveal the phase-related similarity between content-specific representations^7^, we provide evidence for memory-reactivation processes during wakefulness and their reoccurrence during NREM sleep.

**Figure 1.**
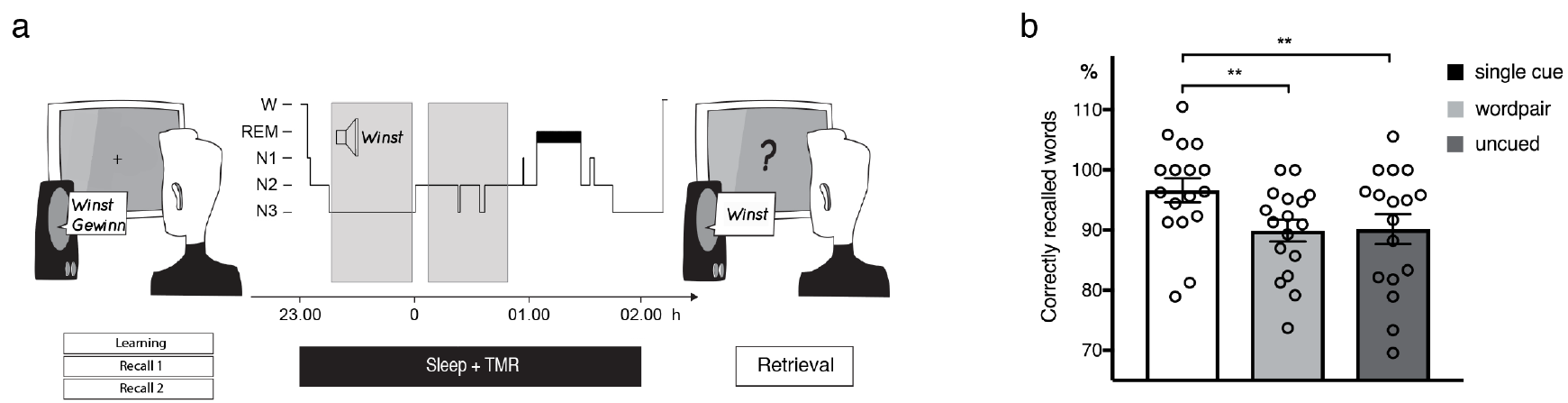
Experimental design and behavioral results. **(a)** Participants performed a vocabulary-learning task in the evening. They learned to associate Dutch words (cues) with German words (targets). After the initial learning phase, a cued recall including feedback was performed (‘recall1’). The cued recall was repeated without feedback (‘recall2’). Subsequently, participants slept for 3 hours. During NREM sleep, 80 Dutch words (40 cued, 40 cued+feedback) were repeatedly presented. Memory performance was assessed in the final retrieval phase after sleep **(b)** Presenting single Dutch word cues during NREM sleep enhanced memory performance as compared to word-pair TMR and uncued words. Retrieval performance is indicated as percentage of recalled words, with performance before sleep set to 100%. Values are mean ± s.e.m. **P<0.01.

In a first step we evaluated, whether we could identify content-specific phase patterns of memory retrieval during wakefulness, indicating recall-related memory reactivation. The degree of phase similarity for retrieving the same memory content (i.e., word) during consecutive recall instances (recall1 / recall2) was assessed using the pairwise phase consistency and contrasted between remembered and non-remembered words (Supplementary Fig. 1 + 2) for frequencies between 3 and 16 Hz.

We found significantly higher phase similarity for remembered words as compared to non-remembered words in the theta range (cluster-randomization: *P* = 0.006, corrected for multiple comparisons), peaking at 5Hz (Fig. 2a+b). The time-course of the phase similarity at 5 Hz displayed an early significant difference between remembered and non-remembered words (cluster-randomization: *P* = 0.008, corrected for multiple comparisons; Fig. 2c). Additional analyses indicated that the phase similarity results were not biased by spectral power (see Supplementary Results). Given our design, it seems unlikely that those results were driven by similarities in auditory stimulation. Still, we tested this possibility by assessing phase-similarity between learning and both recall instances, with the learning data being segmented around the onset of the Dutch words (thus before any association was learned; see Methods). No significant cluster was observed (both *P’s* > 0.3).

**Figure 2.**
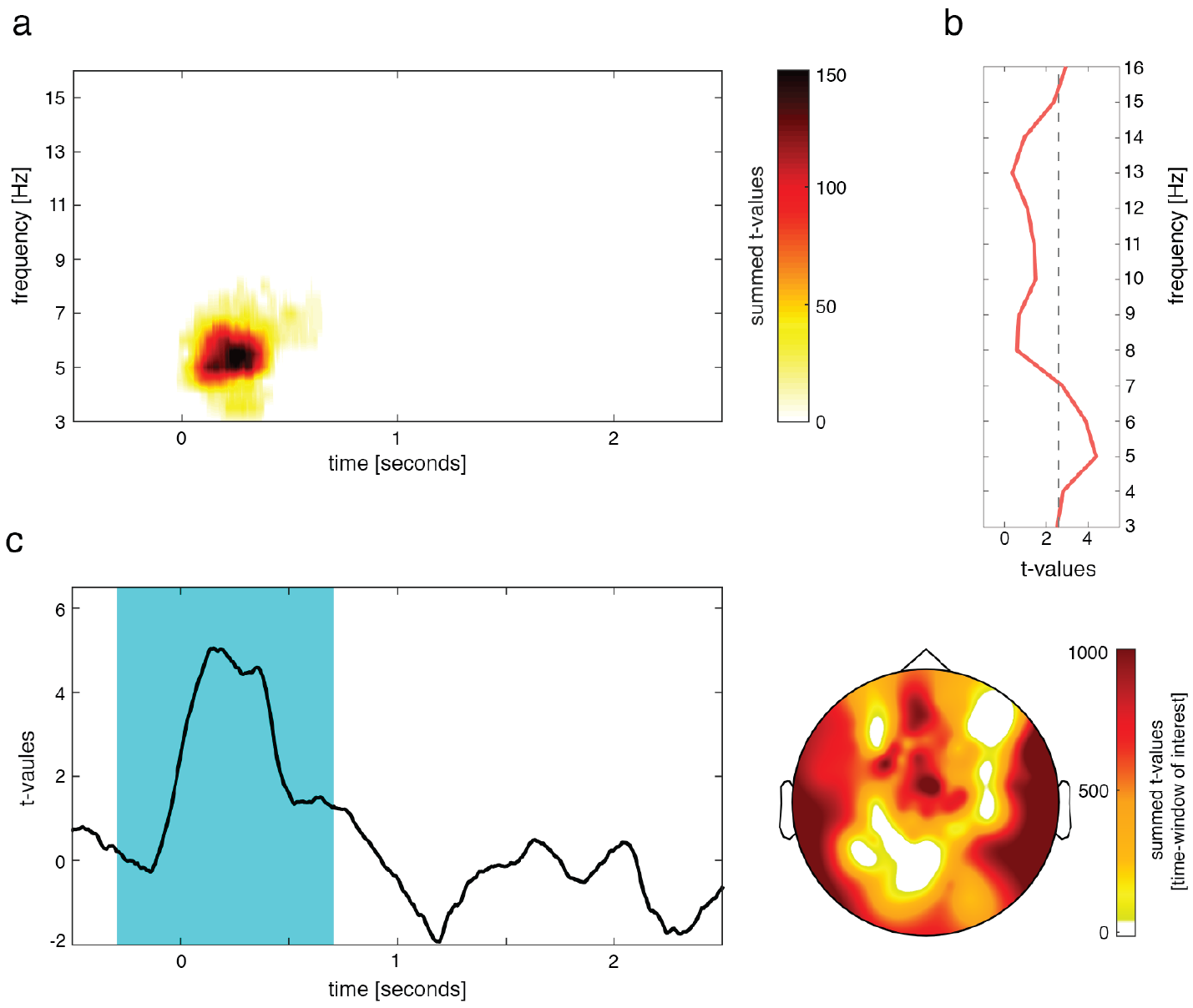
Word specific phase-similarity during wake retrieval. **(a)** Significantly enhanced phase similarity during successful subsequent retrieval was observed early after cue onset (t = 0 sec) in the theta range **(b)** peaking at 5Hz. **(c)** time-course and topography of phase similarity at 5Hz, indicating a rapid reactivation of memory content. The one second time-window around the center of the strongest cluster is highlighted.

The next crucial step was to test whether these content specific features tracked by phase-similarity at 5 Hz would be shared between reactivation processes during wakefulness (‘recall2’) and sleep (TMR). Because memory reactivation during sleep could emerge at any point after TMR cue presentation, phase-similarity between recall2 and TMR was examined with a sliding window approach^7^, using the single-trial phase locking value. Target words remembered after sleep were paired with their equivalent during ‘recall2’ and contrasted against non-remembered words. A one second time-window exhibiting the strongest content-specificity from the pre-sleep retrieval (center: 0.193ms) was used as sliding window. Test statistics on the averaged difference between remembered and non-remembered words revealed the reactivation of recall-related phase-patterns at 5 Hz during TMR during sleep (cluster-randomization: *P* = 0.02, corrected for multiple comparisons) over right temporal electrodes (Fig. 3a). There was no significant difference in spectral power biasing the results (see Supplementary Results). To assess the frequency specificity of the obtained results, the same analysis was performed for 3Hz and 8Hz. No significant effects were found (both *P* > 0.16).

**Figure 3.**
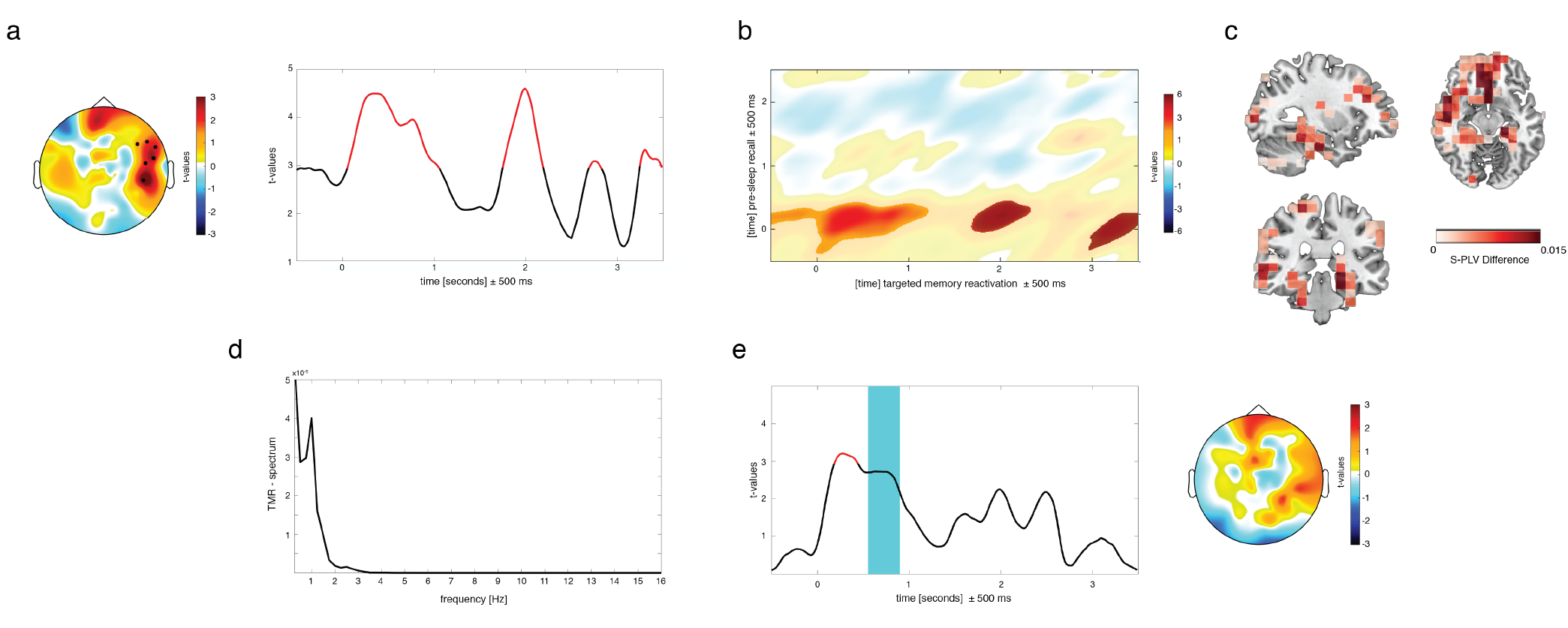
Retrieval-TMR similarity. **(a)** Re-current reactivation of recall-related phase-patterns at 5 Hz during TMR emerged over right temporal electrodes **(b)** Assessing phase-similarity between every time-point of retrieval and TMR confirmed the cycling result pattern **(c)** Source reconstruction of the obtained effects indicating differences in right (para)hippocampal regions and left-frontal areas **(d)** Frequency spectrum of the TMR similarity measures showed a 1 Hz periodicity of reactivation processes. **(e)** In line with behavioral predictions, providing a target stimulus after the TMR-cue blocked associated reactivation processes.

To examine the time-course of the reactivation effect, similarity-measures were averaged across significant electrodes and t-statistics were computed for every time-point. Four distinct reactivation episodes emerged, peaking at 390ms (*t*_16_= 4.49, *P* = 0.0003), 1990ms (*t*_16_ = 4.59, *P* = 0.0002), 2760ms (*t*_16_ = 3.08, *P* = 0.007) and 3310ms (*t*_16_ = 3.31, *P* = 0.004). Importantly, this pattern of results suggests that presenting a memory cue during sleep triggered a repetitive cascade of memory-reactivation, with reactivation processes fluctuating at a frequency of ~1 Hz (Fig.3d). Testing all combinations of recall and TMR time windows revealed that no additional recall episodes were reactivated during TMR (Fig. 3b).

To evaluate the sources of the scalp-level effects, phase similarity was assessed on virtual sensors by applying a DICS beamformer. Source level contrasts revealed differences in right (para)hippocampal regions as well as more widespread difference in left frontal areas, including the frontal gyrus and insula (Fig. 3c).

Our analysis focused on similarity measures between recall and single-cue TMR, because presenting Dutch-German word-pairs during sleep abolished the beneficial effects of TMR on later memory performance^8^. Based on this behavioral outcome, we predicted that providing both the cue and target word during TMR should block functionally relevant memory-reactivation processes. In line with our hypothesis, there was no significant effect when comparing the averaged difference between subsequently remembered and non-remembered words (*P* > 0.17). As the topographical distribution resembled our main results (Fig.3e), the same electrode cluster was used to characterize the time-course. We found an early reactivation episode peaking at 270ms (*t*_16_= 3.2, *P* = 0.005) for word-pair TMR, thus before the onset of the second word. No later episode was observable, indicating that the presentation of a second stimulus may have blocked further memory reactivation.

We show for the first time that memory-related reactivation processes during wakefulness and sleep share the same neural signatures in humans. Specifically, theta oscillations at 5Hz orchestrated the reactivation of memories, irrespective of the physiological state. This result thereby extends previous findings, indicating a role for theta oscillations in mediating communication between the medial temporal lobe and neocortical regions^9^ and by that the cortical reactivation of memories during wake retrieval^10^. Importantly, hippocampal reactivations are also thought to drive consolidation processes during sleep, leading to the integration of newly acquired memories into cortical networks^1^. Consistently, our source level results corroborate this assumption as not only right (para)hippocampal areas, previously associated with successful TMR in humans^11^, but also language-related regions in the left frontal cortex^12^ showed the phase similarity effects.

Moreover, presenting memory cues during sleep triggered a cascade of reactivations. After providing a TMR cue, memory-related reactivation patterns re-emerged autonomously at a frequency of ~1 Hz, which is in line with the assumption that slow oscillatory activity during sleep guides reactivation processes^1,5^. This result also tightly parallels previous work in rodents, which demonstrated that presenting auditory cues during sleep biases the content of associated memory reactivations with maintaining the biasing effect for multiple seconds^13,14^. Importantly, presenting a second stimulus abolished this fluctuation and diminished the beneficial effects of TMR. In sum our results demonstrate the similarity of memory reactivation during wakefulness and sleep, with a cycling and spontaneous re-processing of memories during sleep when triggered by cueing.

## Data and code availability

All data and analysis code are available on reasonable request from the corresponding author.

## Acknowledgements

T.S. is supported by a grant of the Swiss National Science Foundation (SNSF; P2ZHP1_164994). T.St. received funding from the European Union's Horizon 2020 research and innovation programme under grant agreement No 661373. C.F.D.’s research is funded by the Kavli Foundation, the Centre of Excellence scheme of the Research Council of Norway - Centre for Biology of Memory and Centre for Neural Computation, The Egil and Pauline Braathen and Fred Kavli Centre for Cortical Microcircuits, the National Infrastructure scheme of the Research Council of Norway -NORBRAIN, the Netherlands Organisation for Scientific Research (NWO-Vidi 452-12-009; NWO-Gravitation 024-001-006; NWO-MaGW 406-14-114; NWO-MaGW 406-15-291) and the European Research Council (ERC-StG RECONTEXT 261177; ERC-CoG GEOCOG 724836).

B.R. is supported by the SNSF (100014_162388) and the Clinical Research Priority Program (CRPP) “Sleep and Health” from the University of Zurich.

## Contributions

T.S and B.R. conceived the design. T.S. collected the data. T.S. and T.St. analyzed the data. T.S. and T.St. wrote the manuscript. All of the authors discussed the analyses and results and finalized the manuscript.

## Competing financial interests

The authors declare no competing financial interests.

## Corresponding author

Correspondence to: Thomas Schreiner or Tobias Staudigl

## Online Methods

### Participants

The data were taken from Schreiner, Lehmann & Rasch, (2015)^1^. Thus, detailed information about participants, stimuli, task, data acquisition and behavioral results can be found in the original article. From the total of 20 participants (13 female; age: 22.45 ± 2.39) who entered the main EEG analyses in the original study^1^, 3 datasets had to be excluded due to extensive artifacts in the pre-sleep EEG data (recall 1 and 2, for a detailed description please see below).

### Task and Procedure

All participants performed a vocabulary-learning task in the evening (~10pm). The task consisted of 120 Dutch words and their German translation, randomly presented in three rounds. With regards to the first learning round, each trial consisted of a Dutch word, which was succeeded by the German translation. All words were presented via loudspeaker. The trials of the second round (referred to as ‘recall1’) started with the presentation of the Dutch word (cue), followed by a question mark for up to 7 seconds. The participants were asked to vocalize the correct German translation (target) within the 7 seconds (if possible) or to indicate if they were not able to do so. In any case, the correct German translation was presented afterwards. The same cued recall procedure was accomplished in the third round (‘recall2’), except that performance feedback was omitted (for a detailed scheme for recall1 and recall2 see Supplementary Fig.1).

The learning phase was followed by a 3 hours retention interval of sleep. During NREM sleep subsets of the Dutch cue words learned before the retention interval were repeatedly replayed for 90 minutes via loudspeaker, either as single cues (only the Dutch words), 40 as word pair cues (Dutch and German words) and 40 were not replayed at all. During word pair cueing, the presented word-pairs consisted of correct word-pairs just as learned before the retention interval for 8 participants, while 9 participants were presented with newly formed Dutch-German word-pairs during sleep. Importantly opposed to replaying single cues, cueing of word pairs, irrespective of the category (correct, wrong), was associated with a suppression of the beneficial effects of cueing. Thus, we capitalized our main analysis on the single-cue TMR condition, while the word-pair condition served as important control analysis. In all three categories, the relation of remembered and non-remembered word pairs of the last learning trial before sleep was maintained. Hence, all categories comprised the same number of remembered and non-remembered words before sleep. All words were individually and randomly chosen for each participant using an automatic MATLAB algorithm. After sleep, recall of the vocabulary was tested in a final retrieval phase using a cued recall procedure.

### EEG recording and preprocessing

EEG was recorded using a high-density 128-channel Geodesic Sensor Net (Electrical Geodesics, Eugene, OR). Impedances were kept below 50 kΩ. Voltage was sampled at 500Hz and initially referenced to electrode Cz. Offline EEG preprocessing was realized using BrainVision Analyzer software (version 2.0; Brain Products, Gilching, Germany). Data were offline re-referenced to an average-reference. The continuous EEG was epoched into intervals from 1,000ms before until 3,000ms after word onset. Trials with artifacts (e.g., muscle and movement artifacts) were manually removed after visual inspection. Eye blinks and movements of the pre-sleep EEG recordings (recall1 & 2) were corrected using independent component analysis^2^. For each phase (recall1, recall2 and TMR), segments were categorized based on the subjects’ memory performance in the final retrieval phase into later remembered and non-remembered words. All succeeding analyses steps were realized with MATLAB (the MathWorks) using the open-source FieldTrip toolbox^3^.

### Word specific phase similarity during awake recall

To detect content (i.e. word) -specific phase patterns of successful recall during wakefulness, a modified version of the pairwise phase consistency (PPC) was applied^4,5^. In a first step, oscillatory phase was extracted using complex Morlet wavelets of 6 cycles for all frequencies between 1 and 20Hz in steps of 0.5Hz and 1ms, ranging from 1000ms pre-stimulus to 3000ms after stimulus onset.

To compute the pairwise phase consistency, words remembered after ‘recall2’ were paired with their equivalent during ‘recall1’ and contrasted against words not remembered during ‘recall2’ and their equivalent during ‘recall1’. Thereby, the degree of phase similarity between identical words and associated memory content was assessed during consecutive recall instances (recall1 and recall2; for an illustration see supplementary Fig.1). The rationale hereby is that recall processes associated with remembered items should exhibit a higher content-related similarity to each other as compared to non-remembered ones, given the potentially weak/non-existing reactivation of a memory trace in the latter case. For each pair of trials, the cosine of the absolute angular distance was then computed and finally averaged across all (same or different) combinations^4,5^. Thereby a value, representing the average similarity specifically for each set of combinations, was derived for every electrode, frequency and time-bin and subsequently used for statistics. We assessed phase similarity for the same words (identical auditory cue word presentation) presented during ‘recall1’ and ‘recall2’ and contrasted remembered pairs against non-remembered pairs. Therefore, potential confounding influences of similarity in auditory stimulation or mere perceptual processes should be equal and both conditions (remembered and non-remembered pairs) and thus controlled for. To further strengthen this point, we assessed the phase-similarity between learning and both recall instances. Importantly, data from learning was also segmented with regards to the onset of the Dutch words. Since the presentation of the German translations during learning was a delayed by 3 seconds, no memory reactivation could have been made at this point and the recorded EEG activity primarily mirrors the auditory perception and processing of a particular word.

Furthermore, as power differences can bias phase estimation, we further tested whether there was a significant difference in power between conditions. Initially, we estimated oscillatory power for the very same contrasts as utilized in the phase analyses (i.e. oscillatory power for words remembered after ‘recall2’ was subtracted from power values of ‘recall1’ and contrasted against the difference of power-values of words not remembered during ‘recall2’ and their equivalent during ‘recall1’). In addition, we tested potential differences in oscillatory power for remembered versus non-remembered items, independently for recall1 and recall2 (for a detailed overview see Supplementary Fig. 3). Power values were extracted specifically for 5Hz using a complex Morlet wavelet of 6 cycles.

### Word specific phase similarity between recall and TMR

As described above, we initially focused our analysis to the single-cue TMR condition (the word-pairs condition was used as a control analysis, described below). Thus, we tested in a first step whether content specific features of memories would be shared between recall before and single-word TMR during sleep, indicating that TMR during sleep leads to the reactivation of those properties. Target words remembered after cueing during sleep were paired with their equivalent during ‘recall2’ and contrasted against target words not remembered after sleep-cueing and their equivalent during ‘recall2’. Thus, between those pairs of successfully acquired memories a phase similarity measure was calculated and contrasted against words lacking a stable memory trace. Following Michelmann, et al. (2016)^5^, several restrictions were used when applying the phase-based RSA to specify electrodes and frequency of interest and the utilized time windows. We used 5Hz as frequency of interest, given that content-specific phase similarity peaked at this frequency during wake-recall. Hence, oscillatory phase at 5Hz was maximally content-sensitive during recall before sleep. In addition, the time-window at ‘recall 2’ was confined to 1 second, evenly distributed (±500ms) around the peak of the recall-associated phase-similarity measure of 5Hz. Similarly, electrodes were adopted from showing content-specific phase similarity during wake-recall (83 electrodes in total).

To account for the fact that reactivation processes associated with TMR during sleep could happen at any time after cue presentation, a sliding window approach was utilized. Thereby, phase similarity for all trial combinations was determined between each electrode and time point associated with TMR and the 1 second time window derived from the recall before sleep.

We determined phase similarity with the Single-trial Phase Locking Value (S-PLV)^6^, hence the similarity with regards to phase angle differences over time. Again, phase values were extracted using a complex Morlet wavelet of 6 cycles for frequencies between 1 and 20Hz in steps of 0.5Hz. For computational efficiency phase values were down-sampled to 100Hz. Phase similarity was assessed between every pair of remembered items and contrasted against phase similarity between pairs of non-remembered items. The pre-stimulus interval between −500ms and 0ms was used as padding to slide the recall window into TMR episodes.

Next, we explored the time-course of reactivation processes during sleep. To do so the electrodes derived from the 5Hz cluster displaying the significant difference were averaged and subjected to a series of post-hoc *t*-tests between remembered and non-remembered combinations for every time-point of sleep cueing. Subsequently, we repeated the sliding time window analysis using the same electrodes but varying time windows from the pre-sleep recall. This allowed us to evaluate similarity between every time-point of recall and TMR, given an uncertainty of ±500ms. Afterwards the differences of all combinations were averaged across electrodes. Furthermore, two control frequencies were tested to estimate the frequency specificity. Oscillations at 3Hz and 8Hz, which are maximally different in phase to 5Hz (i.e. golden ratio), were utilized as control frequencies.

As TMR during sleep seemed to have triggered reactivation processes in a recurrent fashion, we evaluated whether the similarity measures would fluctuate at a certain frequency (‘TMR-spectrum’). To this end, we performed a spectral analysis of the time-course of the phase similarity differences. We estimated the spectral power for frequencies between 0.25 and 16Hz by multiplying a hanning taper to the Fourier transformation of the whole trial (−0.5 to 3.5 seconds) and evaluated potential peak frequencies.

Furthermore, the same control analyses with regards to learning and oscillatory power as performed for the recall part were conducted. To evaluate whether our similarity measures between recall2 and TMR were influenced by the similarity in perceptual processing, we assessed the phase-similarity between learning and TMR, again with the learning data being segmented with regards to the onset of the Dutch words (thus, before the onset of the translation). To test whether power differences might bias the phase estimation, we investigated oscillatory power for the very same contrasts as utilized in the phase analyses (i.e. oscillatory power for words remembered after ‘recall2’ was subtracted from power values of ‘TMR’ and contrasted against the difference of power-values of words not remembered during ‘recall2’ and their equivalent during ‘TMR’). In addition, we tested potential differences in oscillatory power for remembered versus non-remembered items for TMR (for a detailed overview see Supplementary Fig. 3).

Power values were extracted specifically for 5Hz using a complex Morlet wavelet of 6 cycles. For all control analyses the same time-window and electrode selection was utilized as in the main analysis. Finally, we tested whether presenting an additional stimulus in the word-pair TMR condition might have interfered with on-going reactivation processes as the lack of behavioural effects for this TMR condition suggested such an interpretation^1^. Phase-similarity was assessed between recall and word-pair TMR in the same way as for the main analysis.

### Source estimation

To estimate the sources of the obtained effects a virtual electrode approach, applying the Dynamic Imaging of Coherent Source (DICS) beamforming method^7^, as implemented in FieldTrip, was used. A spatial filter for each specified location (each grid point; 10mm^3^ grid) was computed based on the cross -spectral density, calculated for the frequency of interest (5Hz), using a complex Morlet wavelet (see above), for all trials (common filter approach). As time windows of interest served the 1 second time-window derived from ‘recall2’ and the first second of TMR, given the most extended pattern of phase similarity effects during this period. Electrode locations for the 128-channel Geodesic Sensor Net EEG system were co-registered to the to the surface of a standard MRI template in MNI (Montreal Neurological Institute) space using the nasion and the left and right preauricular as fiducial landmarks without individual digitization. A standard leadfield was computed using the standard boundary element model^8^. The forward model was created using a common dipole grid (10mm^3^ grid) of the grey matter volume (derived from the anatomical automatic labeling atlas^9^) in MNI space, warped onto standard MRI template, leading to 1457 virtual sensors. Subsequently, data analysis was accomplished in the same way on the virtual data as before on sensor level.

## Statistics

### Recall specific phase similarity

Statistical testing of differences in phase similarity between remembered and non-remembered words of ‘recall1’ and ‘recall2’ was accomplished using a cluster-based nonparametric permutation approach^10^. This approach controls the Type I error rate with regard to multiple comparisons (here: time, frequency and space) by clustering neighboring sensor pairs (minimum 2 electrodes), exceeding a critical t-value in the same direction. For all included frequency bins (3-16Hz), paired sampled t-tests were computed for any given electrode and for each time-point (ranging from −0.5 to 2.5 seconds with regard to stimulus onset). Thereby, clusters of contiguous sensors across participants were identified (*P* <0.05, two-tailed). The cluster-level statistic was defined from the sum of the t values of the sensors in a given cluster. Only the cluster with the largest summed value was considered and tested against the permutation distribution (Monte-Carlo method, *P* < 0.05, two-tailed t-test). To estimate the time-course and the topographical distribution of the peak frequency, effects at 5Hz, were tested specifically using the same procedure as above against a one-sided distribution (controlling for multiple comparisons in time and space).

### Phase similarity between recall and TMR

Statistical quantification of the phase similarity between ‘recall2’ and ‘TMR’ contrasting remembered and non-remembered words was again accomplished using a cluster-based nonparametric permutation approach. Initially, the average difference between 0 and 3.5 seconds was tested using paired sampled t-tests (*P* < 0.05, two-tailed, controlling for multiple comparisons across space). To correct for multiple comparisons 500 permutations were drawn and the cluster with the largest summed t-value was tested against the permutation distribution. To quantify the temporal characteristics of the obtained effects, phase similarity measures were averaged across the electrodes within the significant cluster. Paired sampled t-tests were computed for every time-point (*P* < 0.01, two-tailed).

### Cluster specific estimation of memory reactivation between recall and TMR

To statistically test similarity differences between varying time windows from the pre-sleep recall2 and TMR a series of post-hoc t-tests was accomplished within the cluster of significant electrodes. T-values, thresholded against a *P*-value of 0.01 (one-sided) were summed up and 500 permutations were drawn. Following Michelmann and colleagues^5^ a distribution comprising the strongest cluster, the second strongest cluster etc. was formed. Subsequently, the obtained clusters were compared against a random cluster distribution. Specifically, the cluster showing the highest sum of t-values was compared with the distribution of the maximum cluster, while the next cluster was compared to the second strongest cluster etc. Finally, *P*-values were divided by the number of clusters (bonferroni-correction).

